# A thermostable oral SARS-CoV-2 vaccine induces mucosal and protective immunity

**DOI:** 10.1101/2021.09.09.459634

**Authors:** Bertrand Bellier, Alicia Saura, Lucas A. Luján, Cecilia R. Molina, Hugo D. Lujan, David Klatzmann

## Abstract

An ideal protective vaccine against SARS-CoV-2 should not only be effective in preventing disease, but also in preventing virus transmission. It should also be well accepted by the population and have a simple logistic chain. To fulfill these criteria, we developed a thermostable, orally administered vaccine that can induce a robust mucosal neutralizing immune response. We used our platform based on retrovirus-derived enveloped virus-like particles (e-VLPs) harnessed with variable surface proteins (VSPs) from the intestinal parasite *Giardia lamblia*, affording them resistance to degradation and the triggering of robust mucosal cellular and antibody immune responses after oral administration. We made e-VLPs expressing various forms of the SARS-CoV-2 Spike protein (S), with or without membrane protein (M) expression. We found that prime-boost administration of VSP-decorated e-VLPs expressing a pre-fusion stabilized form of S and M triggers robust mucosal responses against SARS-CoV-2 in mice and hamsters, which translate into complete protection from a viral challenge. Moreover, they dramatically boosted the IgA mucosal response of intramuscularly injected vaccines. We conclude that our thermostable orally administered e-VLP vaccine could be a valuable addition to the current arsenal against SARS-CoV-2, in a stand-alone prime-boost vaccination strategy or as a boost for existing vaccines.

## Introduction

Current field observations show that a protective vaccine against SARS-CoV-2 is likely the only means of controlling the pandemic^1–3^. To fulfill this promise, these vaccines should ideally be effective in preventing infection and virus transmission and, importantly, well accepted by the population. In underdeveloped countries, vaccines should also have a simple logistic chain^4,5^.

Regarding efficacy, years of vaccine research have demonstrated that vaccine protective effects rely in large part on systemic neutralizing antibodies, while local cytotoxic T cell responses are for the most part responsible for virus eradication after a productive infection^6^. Moreover, for upper respiratory tract infection, a robust mucosal immunity is likely required to minimize virus transmission^7–9^. Regarding vaccine hesitancy, oral administration would favor acceptance and minimize the risk of adverse events^10^. Regarding logistics, oral administration would also ease mass vaccination, and a thermostable vaccine would ensure a much-simplified logistic chain. Likewise, an optimal vaccine against SARS-CoV-2 should be thermostable, orally administered and able to induce a robust mucosal neutralizing immune response.

To tackle the challenge of producing such a vaccine, we used our platform based on retrovirus-derived enveloped virus-like particles (e-VLPs) that has been developed to generate neutralizing antibody (NAb)^11^. Indeed, these e-VLPs have the same lipid membrane as the cell they derive from. Likewise, virus envelope proteins that VLPs express have the same conformation as they have on the lipid membrane of an infected cell, and on the virus itself. As NAbs are mostly targeted to conformational structures, e-VLPs are thus particularly suitable for NAb induction^12^. We previously showed that such e-VLPs could generate robust NAbs against many viruses, such as influenza, HCV and CMV, in mice, macaques and humans^13–17^. Moreover, exploiting the versatile engineering possibilities for these e-VLPs, we recently showed that they could be harnessed with variable surface proteins (VSPs) from the intestinal parasite *Giardia lamblia*, affording them resistance to degradation and the triggering of robust mucosal cellular and antibody immune responses after oral administration^18,19^.

We used this experience to design and evaluate a thermostable orally administered e-VLP vaccine against SARS-CoV-2. We tested the expression of various forms of the Spike protein (S) with or without SARS-CoV-2 membrane protein (M) expression. We found that e-VLPs expressing a pre-fusion stabilized form of S plus M trigger robust mucosal NAbs against SARS-CoV-2 in mice and hamsters, which translate into complete protection from a viral challenge. We consider that such a vaccine could be part of the arsenal against SARS-CoV-2, in a stand-alone prime-boost vaccination strategy or as a boost for existing vaccines.

## Materials and Methods

### Viruses

SARS-CoV-2 isolates were propagated in Vero E6 cells in Opti-MEM I (Invitrogen, Cat. # 51985091) containing 0.3% bovine serum albumin (BSA) and 1 μg of L-1-tosylamide-2-phenylethyl chloromethyl ketone-treated trypsin per mL at 37 °C.

### Experimental animals

For immunization and challenge, the group sizes were chosen based on previous experience and littermates of the same sex were randomly assigned. The number of animals for each experiment and all procedures followed the protocols approved by the Institutional Committee for Care and Use of Experimental Animals. Six week- or four month-old male and female BALB/c mice were used for initial experiments, 6-month-old female and male Golden Syrian hamsters were used in the immunization studies, and 1-month-old female and male SPF Golden Syrian hamsters were used in the challenge experiments. For challenge experiments, under ketamine−xylazine anesthesia, ten hamsters per group were inoculated with 10^5^ PFU of SARS-CoV-2 (in 100 μL) or PBS (mock) via the intranasal route^20,21^. Baseline body weights were measured before infection and body weight was monitored for 28 days (Lee et al., 2020). No animals were harmed during the collection of blood.

### Neutralization assays

Serum taken from intranasally inoculated hamsters at 14 dpi was tested for viral neutralizing antibody titer by microneutralizing assay in Vero E6 cells. Briefly, dilutions of serum samples (1:50 to 1:10,000) were mixed with 100 TCID50 of SARS-CoV-2 virus and incubated at 37°C for 1 h. The mixture was then added to Vero E6 cells and further incubated at 37°C for 72 h^22^. The neutralizing antibody titer was defined as the highest dilution that inhibits 50% of the cytopathic effect.

### VLP expression plasmids

For pGag, the cDNA sequence encoding the MLV Gag (Uniprot: P0DOG8.1) without the C-terminal Pol sequence was used^19^. For SARS-CoV-2 spike protein variants, the cDNA sequences were cloned in the phCMV expression vector^19^. All plasmids were verified by sequencing as reported. The SARS-CoV-2 spike protein variants derived from the wild type strain Swt (NC_045512, original Wuhan variant) all having the D615G and the 682RRAR-685GSAS (modFurinCS) mutations. Additional mutations were inserted in specific variant: Sst1: K986P/V987P; Sst2: T791C/A879C; Sst3: S884C/A893C; Sst4: G885C/Q913C; Sst5: S884C/Q913C.

### VLP generation, production, purification, and validation

VLPs were produced by transient transfection of either HEK293 cells or HEK293-1267 cells, with pGag, pS and its variants, and pM plasmid DNA, using PEI as transfection reagent. Cells were transfected at 70% confluence in T175 flasks with 70 μg of total DNA per flask at a PEI: DNA mass ratio of 3:1. VLP-containing supernatants were harvested 72 h post-transfection, filtered through a 0.45 μm-pore size membrane, and concentrated 20 x in a centrifugal filter device (Centricon® Plus-70-100 K, Millipore Cat. # UFC710008) and purified by ultracentrifugation through a 20% sucrose cushion in an SW41T Beckman rotor (25,000 rpm, 4 h, at 4 °C). Pellets were resuspended in sterile TNE buffer (50 mM Tris-HCl pH 7.4, 100 mM NaCl, 0.1 mM EDTA). Proteins were measured using the Bradford method. For western blotting, proteins were resolved by 10% SDS–PAGE and transferred onto PVDF membranes before incubation with specific primary antibodies. Alkaline phosphatase-conjugated secondary antibodies were used and were detected by BCIP/NBT substrate^19^.

### Immunizations

Mice and Golden Syrian Hamsters were fasted for 4 h and then orally immunized with two weekly doses of 100 μg of different VLPs. For IM immunization, two weekly doses of 10 μg of different VLPs were administered. Animals from the negative control group (naive) received oral immunizations with vehicle alone.

### Fluid collection

Blood was collected weekly from the retro-orbital sinus of hamsters and serum was separated and stored at −80 °C. BAL was collected through the trachea by injection-aspiration of 1 mL of PBS with protease inhibitors.

### Enzyme-linked immunosorbent assay (ELISA) tests

The levels of IgG and IgA antibodies against spike protein were determined by ELISA by sensitizing the plate with homogenates of killed whole virus produced *in vitro*. Spike was quantified using purified protein (Human coronavirus HCoV-229E Spike Protein (S1+S2 ECD). Sino Biological, Inc. 40605-V08B and Spike Protein (Active Trimer) R&D Cat. # 10549-CV). The following secondary antibodies were used: Mouse anti-Hamster IgG Cocktail, Clone: G94-56, G70-204 (BD Biosciences, Cat. # 554009); Mouse anti-Hamster IgM, Clone: G188-9, (BD Biosciences Cat. # 554035); Hamster Immunoglobulin A (IgA) ELISA Kit (MyBiosources Cat. # MBS029668); Mouse monoclonal (H6) anti-SARS-CoV-2 spike glycoprotein (Abcam Cat. # ab273169).

### Statistical analyses

Prism (GraphPad Software) was used to perform one-way or two-way ANOVA on datasets with Tukey’s multiple comparisons test or the Bonferroni post-test, respectively. All figures show mean ± S.E.M. Statistically significant differences are indicated in each graph as ^*^*p*< 0.05, ^**^*p*< 0.01 and ^***^*p*< 0.001 and ns = not significant.

## Results

### Designing and selecting the immunogens

Initially, the spike protein S of SARS-CoV-2 was evaluated and several variants for stabilization of the receptor-binding domain (RBD) and stabilization in the pre-fusion state were designed (Fig.1A). Point mutations, Cys-molecular clamps, furin-cleavage site elimination, and Proline (Pro) substitutions^23,24^ were generated and cloned. Spike protein variants that conserved their own cytoplasmic tail (CT), although it can be advantageous to swap it for the CT of VSV-G to improve pseudotyping onto eVLPs, or that were modified to delete its ER retention signal were also designed. Then, those VLPs were produced and validated for the correct composition as described^25^. VSP-pseudotyped VLPs were orally administered to BALB/c mice and the level of serum IgGs was determined by ELISA (Fig. 2).

**Figure 1.**
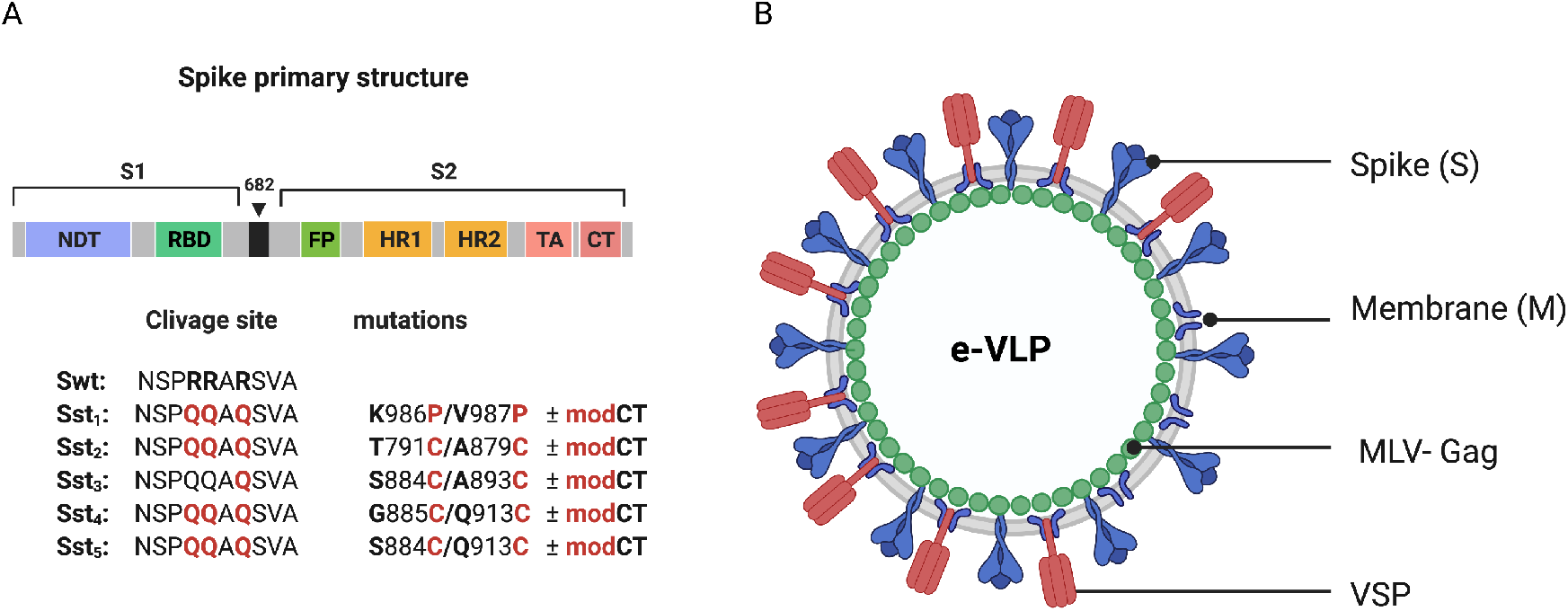
Structure and organization of the SARS-CoV-2 vaccine. (A) Linear diagram of the sequence and structure elements of the SARS-CoV-2 spike proteins used as immunogens. Structural elements include the S1 and S2 ectodomains derived from the original Wuhan variant (Swt) in which specific mutation were inserted. The native furin cleveage site was mutated (RRAR → QQAQ) in all variants (Sst_1_-Sst_5_) to be protease resistant. Specific subsitution (in red) and respective position were indicated. The spike variants with a CT modified to abolish ER retention (modCT) were also generated. (B) SARS-CoV-2 eVLP structure. Native or mutated form of the SARS-CoV-2 spike (S) were pseudotyped on VLP formed with the viral matrix protein MLV-Gag in association or not with the SARS-CoV-2 M proteins and the VSP from the intestinal parasite Giardia lamblia.

**Figure 2.**
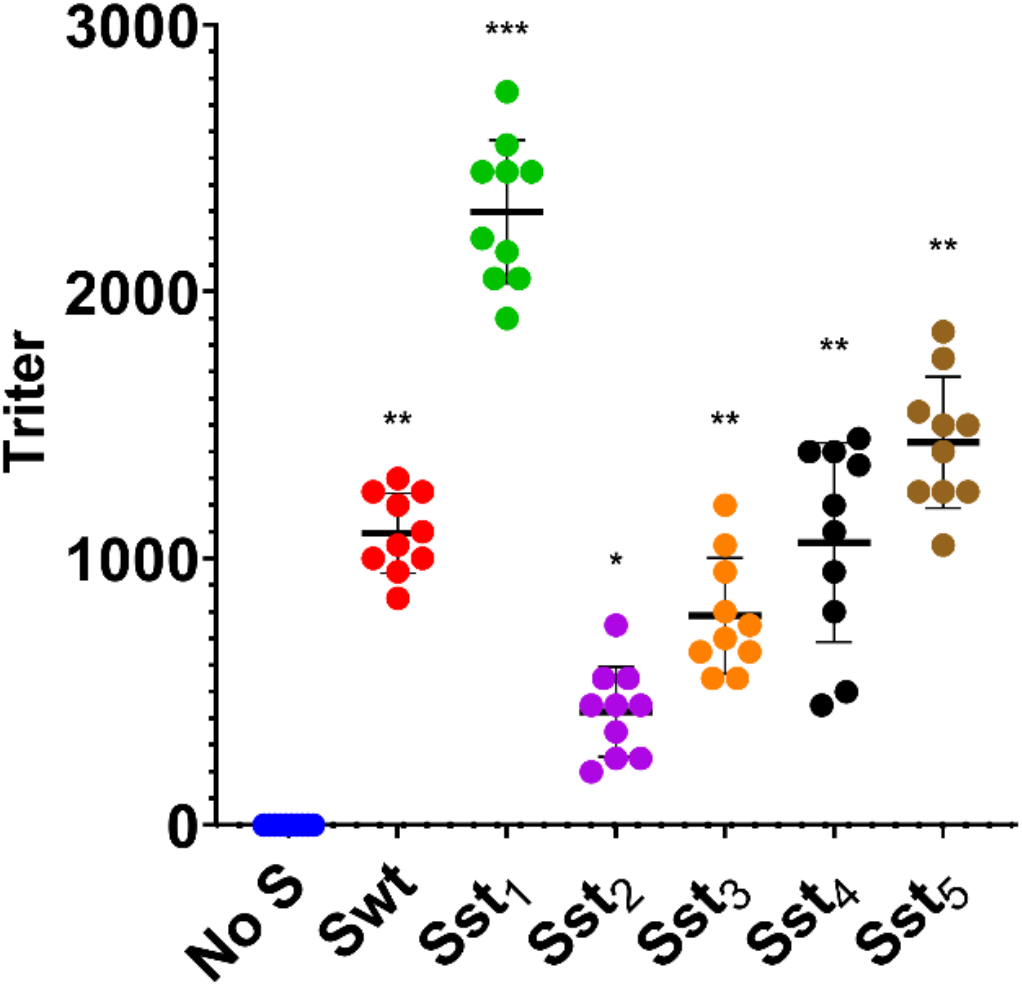
Immunogenicity of different variants of SARS-CoV-2 spike-e-VLPs in mice after oral immunization. VSP-pseudotryped e-VLPs displaying the following SARS-CoV-2 spike protein variants (Sst1, Sst2, Sst3, Sst4, Sst5) or wilt type sequence (Swt) were produced and used for oral immunization as described in Methods. No S means e-VLPs without spike. Values represent the IgG titer in blood of each animal and the horizontal line indicates the mean value. Stabilized Spike 1 (Sst1) presented the higher titers and was selected for subsequent experiments.

Notably, spike variants lacking the furin-cleaved site and with the two Pro substitutions (Sst1), as reported by Wrapp *et al*. (2020)^26^, including the D614G mutation^27,28^, was the most efficient in eliciting a high level of antibodies. Identical experiments were performed in ob/ob mice (JAX™ Mice Strain), db/db mice (JAX™ Mice Strain) and aged mice (>4 month-old) of both sexes and yielded similar results (not shown), indicating that antibody production after oral immunization with e-VLPs does not vary according to underlying condition, sex or age of the mice. Therefore, Sst1, the most effective variant of S, was selected for further studies.

Given that glycosylation of spike would be important for its appropriate conformation since there are numerous glycosylated sites near the RDB ^29,30^, glycosylation of S should likely influence the generation of efficient NAbs. As the C-terminal CT of the SARS-CoV-2 spike is important in proper glycosylation, spike with a CT modified to abolish ER retention (SmodCT) was generated. In addition, the envelope membrane protein M is known to retain S at the ER for improvement of the first steps of glycosylation and, subsequently, remains attached to the CT of S during their journey throughout the Golgi apparatus, where final glycosylation is accomplished^31^. Consequently, e-VLPs with protein M of SARS-CoV-2 were also tested (Fig. 1B). Although that was the main reason for including M in the e-VLPs, subsequent reports showed a specific T cell response to several epitopes of this protein in patients who recovered from COVID-19^32–34^. Thus, incorporation of M in the envelope of the e-VLPs could not only benefit proper glycosylation of S, but also the production of a stronger cellular response to the virus. Therefore, e-VLPs with or without M and with and without a modified CT were generated. Finally, all these e-VLPs were generated with or without the incorporation onto the VLP surface of a VSP derived from *Giardia*^25,35^.

### e-VLP immunogenicity after intramuscular injections

e-VLPs administered intramuscularly (i.m.) to hamsters, whether or not decorated with VSPs, induced high levels of IgG (Fig. 3, left) and moderate level of IgA (Fig. 3, right) in serum, validating these immunogens. Noteworthy, the presence of the *Giardia* VSP on the e-VLPs promoted higher levels of antibodies than the plain e-VLP, highlighting the adjuvant effect of the VSPs^25,35^.

**Figure 3.**
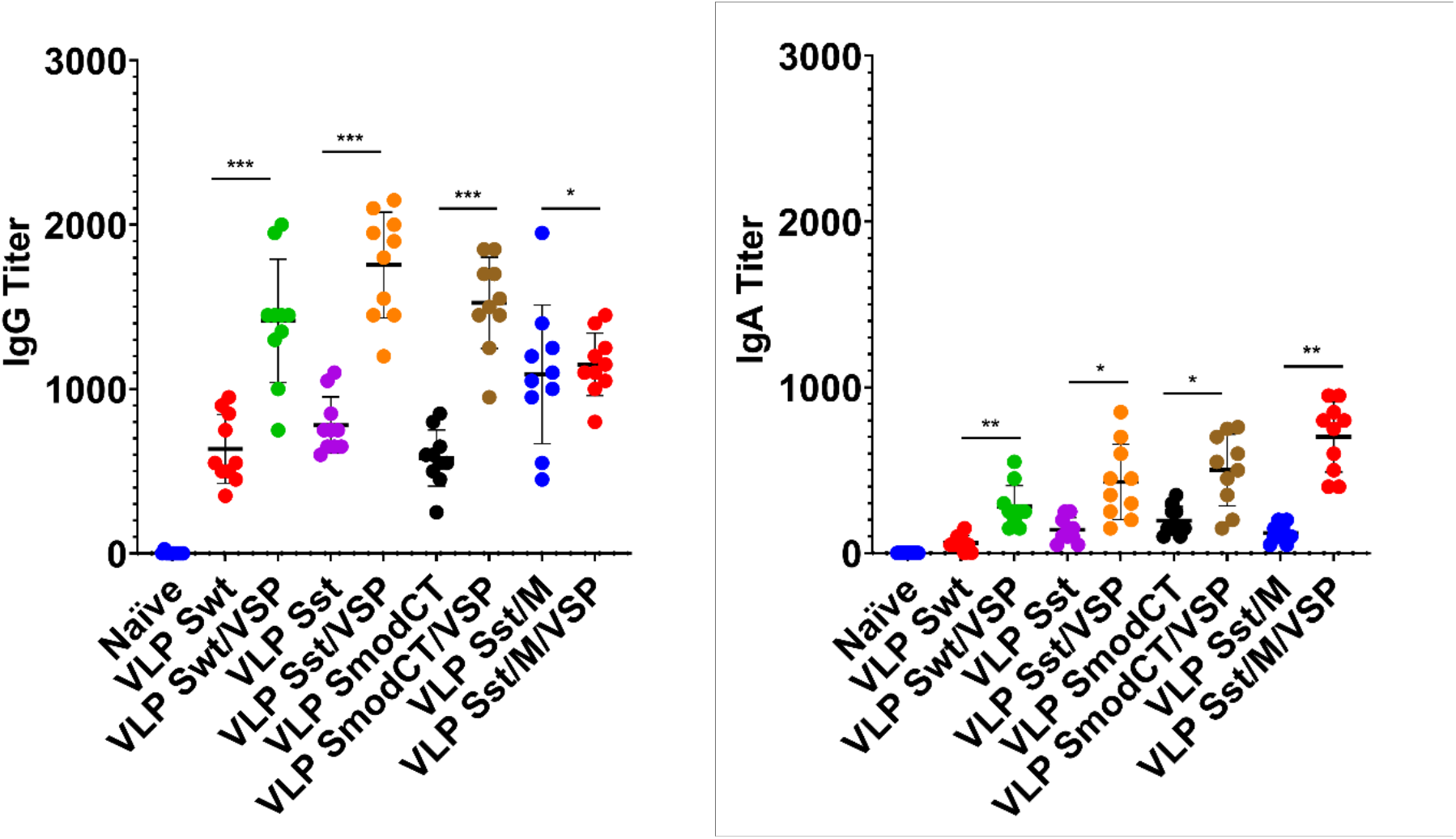
Serum antibody responses to intramuscular administration of different vaccine formulations in hamsters. Serum IgG (left) and serum IgA (right) titers of hamsters unvaccinated (naïve) or vaccinated intramuscularly with different formulations. Wild-type spike (Swt), stabilized S as in Fig. 1 (Sst) and the same in which the cytoplasmic tail was modified (SmodCT) were used to pseudotype e-VLPs including or not SARS-CoV-2 M protein and *Giardia* VSPs. Data were analyzed by one-way ANOVA and Sidak’s comparison test. Values represent the mean ± s.e.m. \**p*<0.05; \*\**p*<0.01; \*\*\**p*<0.001; ns, not significant.

The levels of serum IgG after immunization with Sst1 were higher than with the wild-type spike, and the addition of M further increased the IgG response. The levels of serum IgGs were also higher than those of IgA were, as expected. However, these values are comparable to if not higher than those obtained after immunization with other vaccine formulations^36,37^ or those found in plasma from convalescent patients^38,39^.

Altogether, these results clearly show that plain e-VLPs are already good immunogens when administered by injection in the absence of any adjuvant, highlighting that VLPs are structures well recognized by the immune system. These responses are strongly increased when VSPs were present on e-VLPs according to the VSPs’ intrinsic TLR4-dependent adjuvant properties^25,35^.

### e-VLP immunogenicity after oral administration

Oral administration of the same validated immunogens showed that the absence of the *Giardia* VSP decorating the different e-VLPs led to no detectable immune response, most likely due to destruction of the VLPs in the upper small intestine (Fig. 4). Additionally, the modification of the CT of S appeared detrimental in inducing either serum IgG (Fig. 4 left) or IgA (Fig. 4 right) as compared with S having the wild-type CT. However, serum IgG and IgA titers were augmented when M was incorporated into the VSP-e-VLPs (Fig. 4). Noteworthy, the serum IgA induced by the VSP-decorated e-VLPs were higher after oral administration (Fig. 4 right) than after i.m. injections (Fig. 3 right).

**Figure 4.**
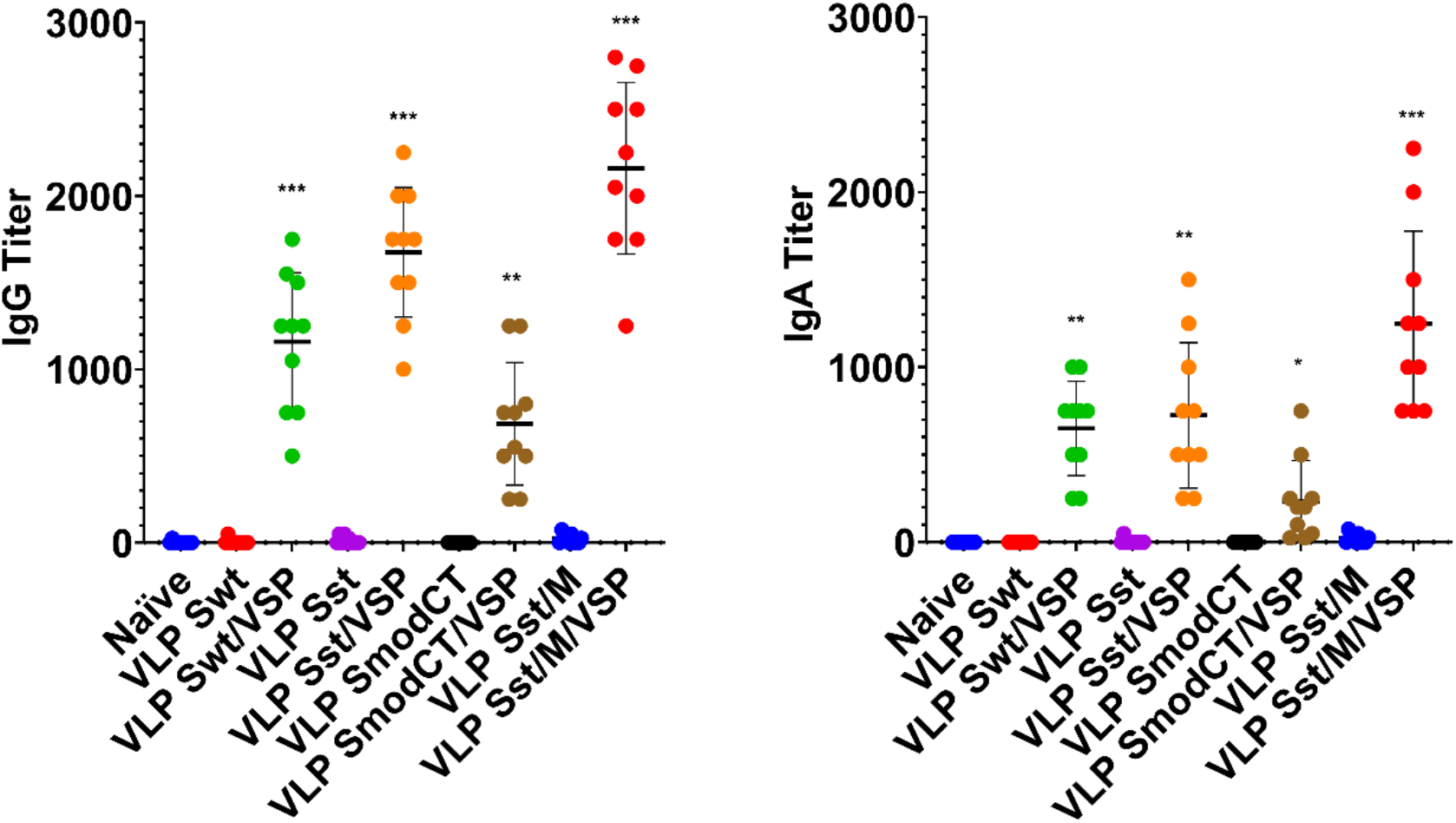
Serum antibody responses to oral administration of different vaccine formulations in hamsters. Serum IgG (left) and serum IgA (right) titers of hamsters unvaccinated (naïve) or vaccinated orally with different formulations. Wild-type spike (Swt), stabilized S as in Fig. 1 (Sst) and the same in which the cytoplasmic tail was modified (SmodCT) were used to pseudotype e-VLPs including or not SARS-CoV-2 M protein and *Giardia* VSPs. Data were analyzed by one-way ANOVA and Sidak’s comparison test. Values represent the mean ± s.e.m. \**p*<0.05; \*\**p*<0.01; \*\*\**p*<0.001; ns, not significant.

Altogether, these results show that the VSPs are essential for oral immunization with VLPs and the immunogenicity of orally administered e-VLPs was strictly dependent on the presence of the VSP on their surface, highlighting the efficiency of this route of immunization, which provides slightly higher Ig responses than after i.m. injection of the same immunogens.

### e-VLP-induced mucosal immunity

When the presence of IgA was analyzed in bronchoalveolar lavages (BAL) of animals immunized orally or by i.m. injection, it was noticed that intramuscularly immunized animals have a consistently low level of IgA titers as compared with those administered orally, in which only the e-VLP version containing VSP, M, and stabilized S showed high titers of IgA (Fig. 5). Again, these results confirm (i) the high immunogenicity of e-VLP formulations, (ii) the crucial role of VSPs in protection of e-VLPs, and (iii) the higher efficiency of oral administration in inducing mucosal IgA.

**Figure 5.**
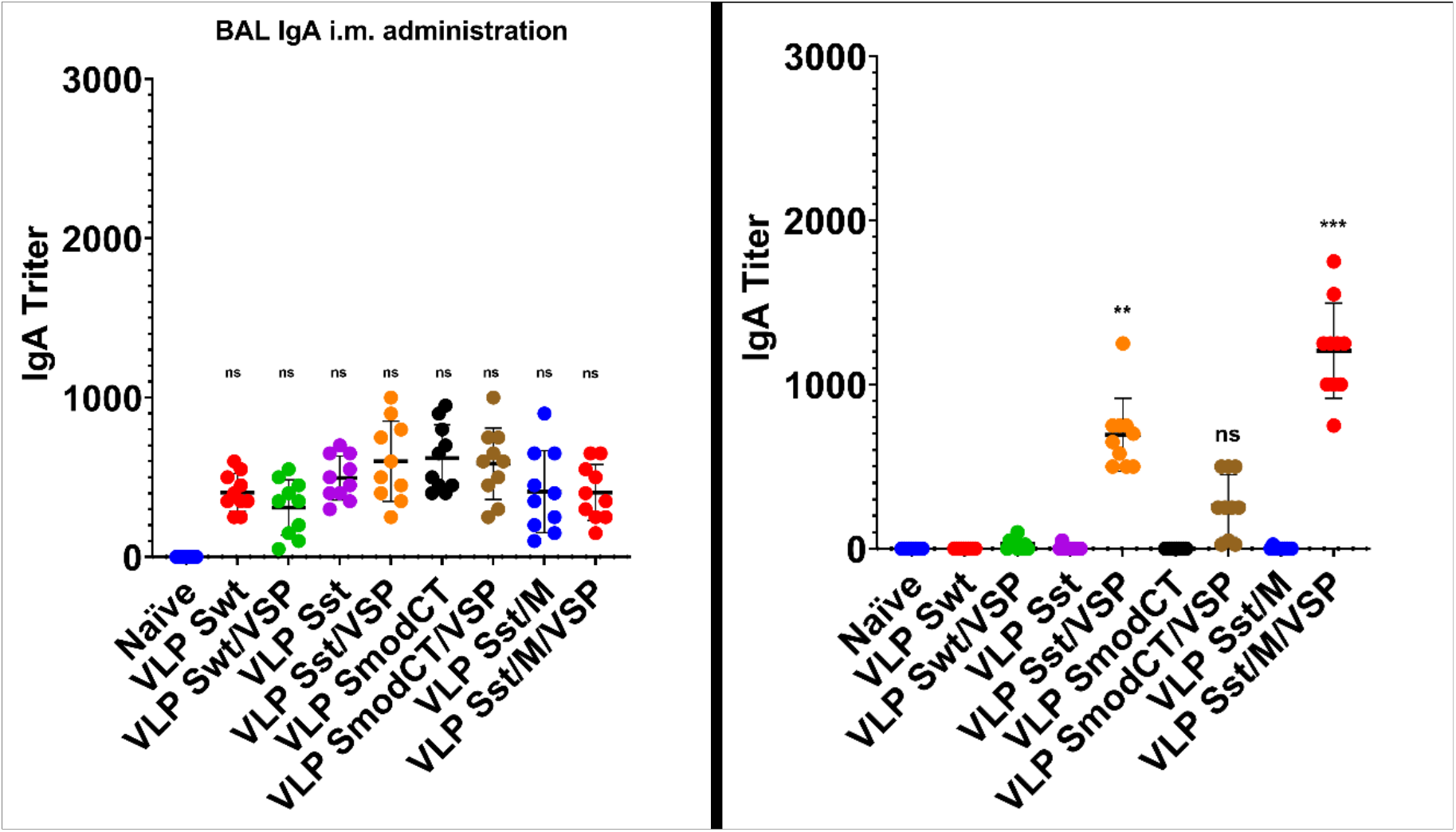
Bronchoalveolar lavage IgA responses. Bronchoalveolar lavage IgA titers of hamsters unvaccinated (naïve) or vaccinated intramuscularly (left) or orally (right) with different formulations. Wild-type spike (Swt), stabilized S as in Fig. 1 (Sst) and the same in which the cytoplasmic tail was modified (SmodCT) were used to pseudotype e-VLPs including or not SARS-CoV-2 M protein and *Giardia* VSPs. Data were analyzed by one-way ANOVA and Sidak’s comparison test. Values represent the mean ± s.e.m. \**p*<0.05; \*\**p*<0.01; \*\*\**p*<0.001; ns, not significant.

### e-VLP-induced neutralizing antibodies

We then compared the best e-VLPs expressing the Sst1 spike and M proteins with the e-VLPs expressing a wild-type spike for the generation of neutralizing antibodies. Interestingly, there was no NAb generated after i.m. injection when the VSP was not present on the VLPs (Fig. 6), highlighting its adjuvant effect. With the VSPs present, both wild-type and stabilized S was able to generate NAbs, as observed for the different commercial vaccines already being administered to humans^1,40–42^. Remarkably, the titer of NAbs generated after oral administration are equivalent to those generated after i.m. injections (Fig. 6), highlighting the efficiency of VSP-e-VLPs as immunogens.

**Figure 6.**
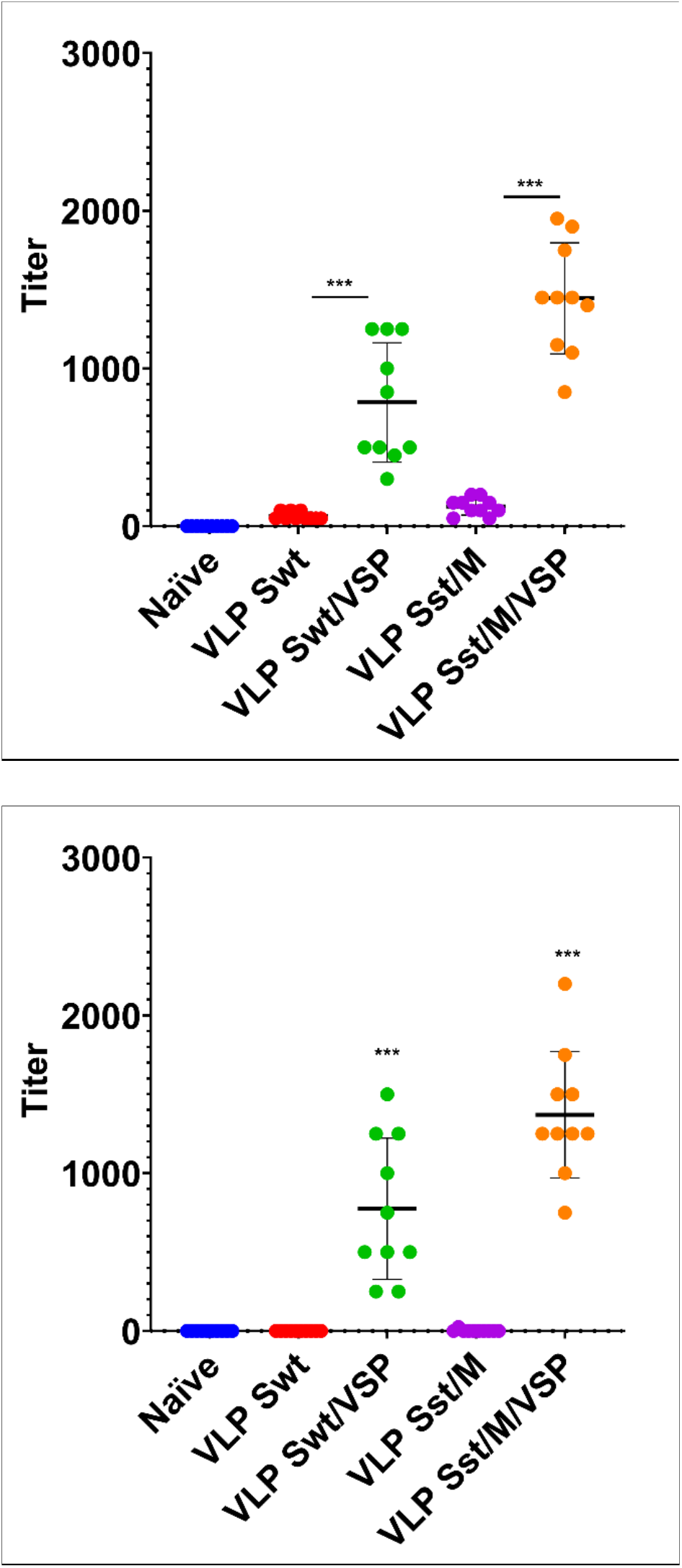
Neutralizing antibodies against SARS-CoV-2 entry. Neutralizing antibody titers of intramuscularly (left) and orally (right) vaccinated animals with selected e-VLP formulations and control animals (naïve). Data between e-VLPs pseudotyped or not with *Giardia* VSPs were analyzed by one-way ANOVA and Sidak’s comparison test. Values represent the mean ± s.e.m. \**p*<0.05; \*\**p*<0.01; \*\*\**p*<0.001; ns, not significant.

### e-VLP as booster immunization

Given the incomplete level of protection afforded by some vaccines and the constant emergence of new viral variants, these orally administered immunogens could possibly be good boosts for existing vaccines. For these reasons, an oral boost was applied to animals previously vaccinated by i.m. injections. In these animals, a third dose of the oral formulation containing VSP, stabilized S and M induced a major increase in the levels of IgA in BAL as compared to those that only received two doses intramuscularly (Fig. 7).

**Figure 7.**
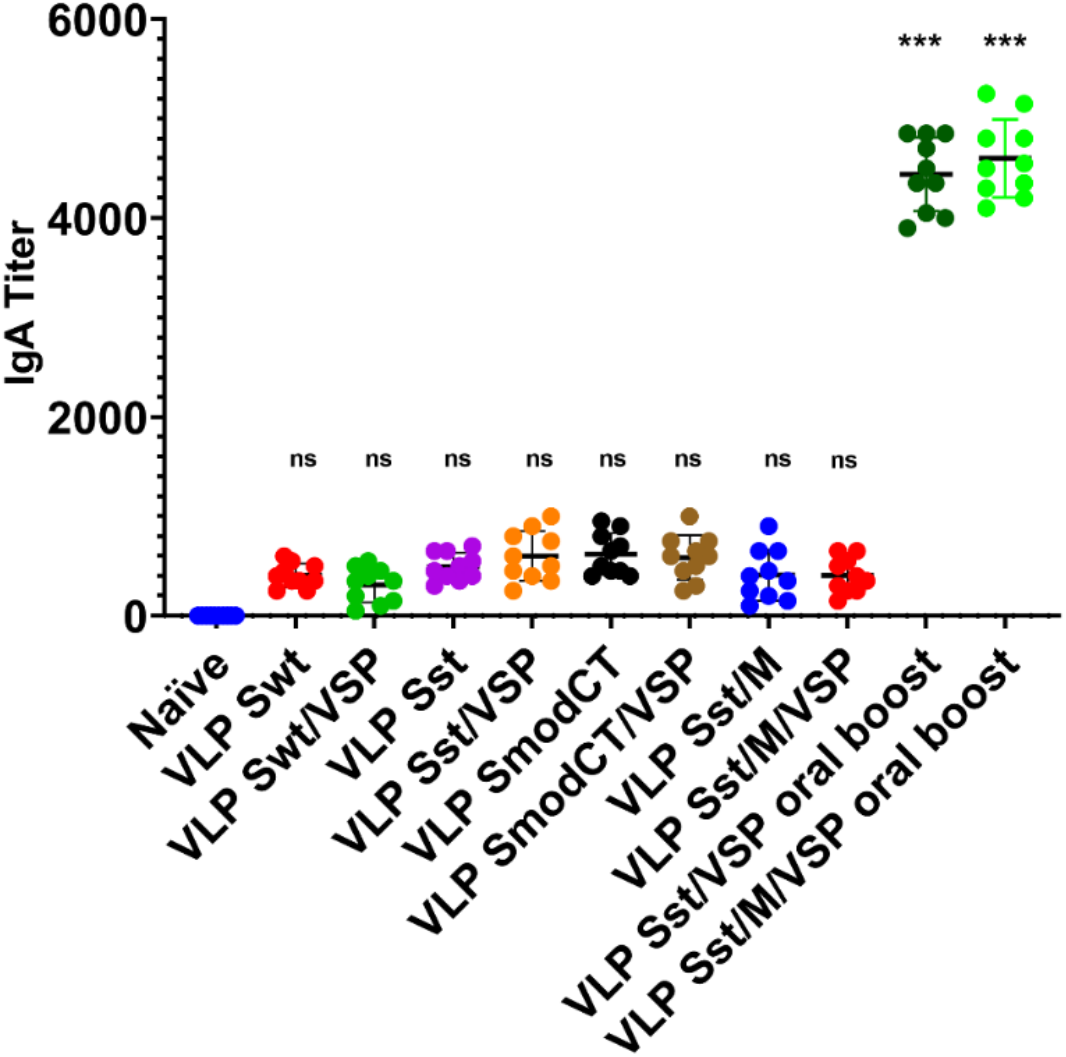
Bronchoalveolar IgA after an oral boost of intramuscularly vaccinated hamsters. Same as Fig. 4a but including an extra oral boost with formulation including stabilized spike and VSP in the presence or absence of SARS-CoV-2 M protein. Data were analyzed by one-way ANOVA and Sidak’s comparison test. Values represent the mean ± S.E.M. \**p*<0.05; \*\**p*<0.01; \*\*\**p*<0.001; ns, not significant.

### Oral vaccination with VSP-e-VLPs protects hamsters from a challenge with SARS-CoV-2

Animals immunized with e-VLPs in which stabilized S and M were present on the particles with or without VSP pseudotyping were challenged with SARS-CoV-2 and the clinical response of the hamsters was determined by monitoring their weight^21^. Control animals lost weight during the two weeks following the viral challenge (Fig. 8) and then recovered, as reported for experimental infections in hamsters^20,21^. Hamsters that were only immunized intramuscularly were not fully protected as they had only a slightly lower weight loss. In contrast, oral immunization with VSP-e-VLPs fully prevented weight loss, whether or not the M protein was present, and similarly to animals that were immunized by injection first and then boosted orally (Fig. 8).

**Figure 8.**
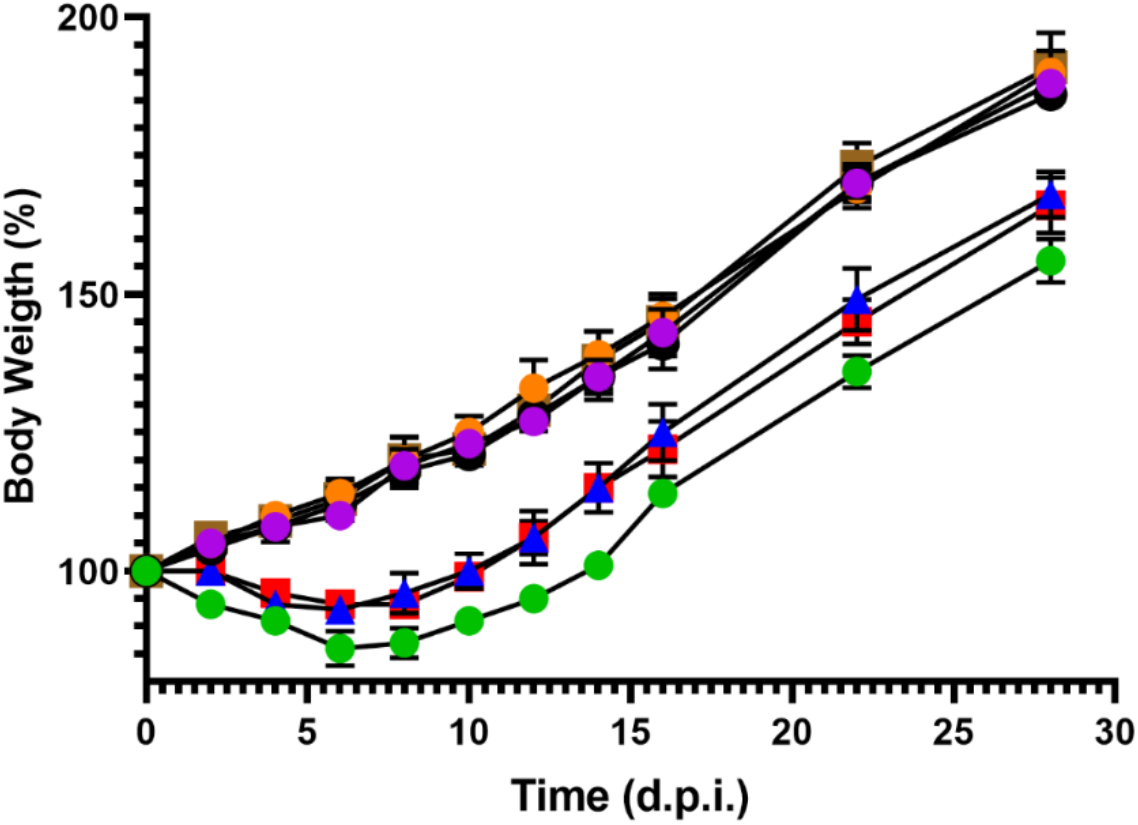
Vaccinated hamsters challenged with SARS-CoV-2. Hamsters either orally or intramuscularly vaccinated were challenged intranasally with purified SARS-CoV-2 virus. Every two days, their weight and general status were monitored and recorded. Unvaccinated animals (green circles); Hamsters orally vaccinated with e-VLPs expressing VSP and stabilized S with the addition or not of the M protein (purple and yellow circles respectively); Intramuscularly vaccinated animals with the same formulations (blue triangles, red squares); oral boost of i.m. vaccinated animals (black circles, brown squares). Values represent the mean of three independent determinations made 1 h apart ± SD.

## Discussion

The differences observed between the same formulations administered either orally or intramuscularly in these animals suggest that although the oral route is expected to show a higher degree of variation among animals, this was not the case. This could be explained by the type of generated Igs. Notably, considering the i.m. administration was done in the absence of any added adjuvant, the high immunogenicity of VSP-e-VLPs can be explained by the adjuvant properties of the VSPs, which have been demonstrated to activate TLR-4^19^. The immunogenicity of the e-VLPs lacking VSPs may mainly rely on the particulate nature of VLPs and the repetitive exposure of the antigen on their surface, even TLR-signaling as been described^43^.

Our results first show that it is possible to co-express SARS-CoV-2 envelope proteins together with *Giardia* VSPs on e-VLPs to generate mucosal Igs and NAbs against SARS-CoV-2 after oral administration. While plain e-VLPs did not generate any Ab responses, VSP-decorated e-VLPs (VSP-e-VLPs) generated Ab responses in the range of, if not higher than, the response to i.m. administration. This is remarkable as it indicates that the SARS-CoV-2 Env proteins are well protected from degradation by VSPs as they preserve their proper conformation, which is needed for NAb production. Thus, this extends our previous results with e-VLP expressing HA of influenza^19^, demonstrating the versatility of the VSP-e-VLP platform. Actually, the dual properties of VSPs were confirmed: they not only afford protection from degradation, but also have a potent adjuvant effect. Indeed, when vaccines are administered i.m., VSP-e-VLP always led to higher titers of antibodies than their plain e-VLPs. Of note, a SARS-CoV-2 VLP based on our platform technology^15^ was independently reported to generate a good NAb response after i.m. administration, but with no reports of IgA at mucosal sites.

Besides its ease, oral administration is known for also having the advantage of triggering better mucosal immunity. This is indeed the case here, with high levels of plasma but also bronchoalveolar lavage IgA detectable only after oral administration. This is an obvious advantage for a vaccine against SARS-CoV-2, as it should reduce viral transmission. In this line, SARS-CoV-2 was still detected in BAL of i.m. vaccinated macaques that otherwise appeared protected from infection. Whether a better mucosal response, as afforded by VSP-e-VLPs, will completely sterilize challenged macaques requires further investigation.

We have not tested the specific T cell response in this study. However, it is known that e-VLPs do induce robust cellular responses; indeed, using VSP-HA-VLPs, a strong cytotoxic T lymphocyte response was generated that was able to kill HA-expressing tumor cells^25^. Moreover, the IgG and IgA responses here are notoriously T cell-dependent and the good antibody response thus attests to a good T cell response^44^. In this line, we previously showed that the fusion of a viral peptide to Gag, the retroviral protein precursor that drives the formation and release of the viral particle/VLPs, produces additional strong T cell responses against this peptide^12^. The fusion to Gag of large fragments or the SARS-CoV-2 N structural protein, or a stretch of immunodominant and/or conserved peptides, would be a mean to further enhance the immunogenicity of VSP-e-VLPs and enhance protection against variants.

SARS-CoV-2 e-VLPs and VSP-e-VLPs could be used as a stand-alone vaccine, likely with a prime-boost scheme of administration. VSP-e-VLPs are thermostable ^19^, retaining their properties at room temperature and tolerating several freeze-thaw cycles, and could thus be particularly advantageous for vaccination in countries where refrigeration of vaccine supplies is problematic. VSP-e-VLPs could also be used as a boost for other vaccine designs. In this regard, it is still unknown how long the protection afforded by the currently used vaccines will last. The follow-up of infected patients indicates that, at least for some patients, the persistence of NAbs and the duration of protection might last a few months^45^. These findings, plus the advent of viral variants, make it likely that the global population will need to boost the immune response of vaccinees regularly. For some vaccine designs, and particularly those based on adenoviral vectors, the re-administration of the same vector might not be very efficient due to the immune response generated against the vector. For these, a boost with VSP-e-VLPs might be particularly interesting. For other vaccine designs, and especially if repeated administrations are needed over the years, an orally administered vaccine might be more acceptable.

The SARS-CoV-2 pandemic calls for vaccination of very large groups of people. This requires a suitable production of vaccine with an excellent safety profile. Noteworthy, we contributed to the design of an anti-CMV e-VLP vaccine based on our e-VLP platform that has already been used in patients, demonstrating scalable GMP production and an excellent safety profile^15,17,46^.

Altogether, given the specific issues of each vaccine design (thermostability, side effects, lack of mucosal immunity induction, immunogenicity against the vector, among other benefits), the availability of multiple vaccines against SARS-CoV-2 improves our chances of controlling the pandemic. In this regard, a thermostable orally administered e-VLP vaccine will be a valuable addition to the current arsenal against this virus.

## Acknowledgments

This work was supported by grants from FONCYT (PICT-E 0234, and PICT-2116), CONICET (D4408), and UCC (80020150200144CC) of Argentina, a Georg Forster Award of the Alexander von Humboldt Foundation of Germany to H.D.L., and recurrent funding by Sorbonne University and INSERM to DK.

## Author contributions

HDL & DK conceived the project. BB, HDL & DK designed the experiments, and supervised students and technicians. AS, LL, CM performed experiments. BB, HDL and DK wrote the paper.

## Competing interests

HL and DK are inventors of a patent application claiming orally administered vaccines against coronaviruses that is owned by their public institutions.

## Notes

### Competing Interest Statement

The authors have declared no competing interest.

